# Intermolecular interactions between cysteine and aromatic amino acids with phenyl moiety in the DNA-binding domain of heat shock factor 1 regulate thermal stress-induced trimerization

**DOI:** 10.1101/2023.11.21.568196

**Authors:** Chang-Ju Lee, Bo-Hee Choi, So-Sun-Kim, David Nahm-Joon Kim, Jeong-Mo Choi, Young-Shang Park, Jang-Su Park

## Abstract

In this study, we investigated the trimerization mechanism and structure of heat shock factor 1 (HSF1) in humans, goldfish, and walleye pollock at various temperatures. The trimerization of HSF1s were confirmed using western blotting using their respective antibodies. First, we examined the HSF1 DNA-binding domains of human (Homo sapiens), goldfish (Carassius auratus), and walleye pollock (Gadus chalcogrammus) by mutating key residues (36 and 103) that are thought to directly affect trimer generation. Humans, goldfish, and walleye pollock contain cysteine at residue 36, but cysteine (C), tyrosine (Y), and phenylalanine (F) at residue 103. Also, the trimer formation temperature of each species was found to be 42, 37, and 20 °C, respectively. In the mutation experiment, trimerization formed at 42 °C when residue 103 was C, at 37 °C it was Y, and at 20 °C it was F, regardless of the species. In addition, it was confirmed that when residue 103 of the three species was mutated to alanine (A), trimer was not formed. This suggest that, in addition to the previously identified C-C disulfide bonds in humans, C forms a trimer with a new type of bond with aromatic ring residues such as Y and F. Thus, HSF1 trimer formation temperature reveals the trimer creation mechanism through the fact that goldfish can have C-Y bonds at 37 °C, and walleye pollock can have C-F bonds at 20 °C. This study suggests that the trimer formation temperature and mechanism of HSF1 are regulated by the amino acid at residue 103.

## 1. Introduction

In recent decades, the rise in water temperature due to global warming has caused many problems in eco systems (Poff et al., 2002). In particular, global warming is expected to increase the water temperature by about 1–4 °C by 2100, and this increase in water temperature induces acute stress reactions in fish in the sea (Alfonso et al., 2021; Barbarossa et al., 2021). The populations of Coldwater fish such as walleye pollock (Gadus chalcogrammus) have already shown a rapid decline in many seas, including along the North Atlantic (Brander, 1997). Also harvest rate of walleye pollock in South Korea are also declining significantly (Lee and Kim, 2010). In addition, because of the increased water temperature, directly or indirectly, aquatic organisms are affected by survival, growth, migration of habitats, population decline, and sexual differentiation (Devlin, 2002; Salinger and Anderson, 2006; Kim et al., 2020). When organisms such as fish are subjected to thermal stress, their cells are exposed to oxidative environments.^1^ And they undergo changes that misfolded or damaged proteins by stress. However, all cells can recover from external stress, including changes in temperature ^2–4^. If misfolded or damaged proteins are present because of temperature changes, some genes encode thermal heat shock proteins (HSPs).^5–6^ HSPs are synthesized in response to stress and are important for preventing proteins from folding incorrectly. Exposure to stress can activate the thermal shock gene and heat shock factor 1 (HSF1) and increase HSP expression. HSPs then help the target protein fold back into its normal form.^7–8^ The transcriptional activation of this protein is regulated by heat shock factors (HSFs), which bind to heat shock elements (HSEs) to mediate the transcription of heat shock genes ^6^. There are four types of HSFs in order of discovery, each of which has a variety of roles, and HSF1 is the representative HSF that responds to temperature stress.^9–11^ Under normal conditions, HSF1 is present inside cells as a monomer. However, when cells are exposed to stressful environments, trimers that participate in the thermal shock response are formed.^12–14^ Previous studies have shown that the trimerization of human HSF1s (hHSF1) occurs at 42 °C when the two cysteines in the DNA-binding domain form a disulfide bond. These two cysteines are located at the amino acid residues 36 and 103 (C36 and C103, respectively). ^15,12^ Additional research has shown that the proximity of aromatic acids is required for the formation of a disulfide bridge between the two cysteines. Without aromatic amino acids such as phenylalanine (F) and tyrosine (Y) around residues 36 and 103, trimerization via the formation of a disulfide bridge does not occur. ^16^ Compared to hHSF1, little is known about the structure and formation of HSF1 and its trimerization in different species of fish. Previous studies conducted in our lab have shown that the goldfish, Carassius auratus, which is a representative example of a temperate fish, has an HSF1 trimerization temperature of 37 °C. However, the mechanism of how goldfish HSF1 forms trimers remain unclear. The structures of hHSF1 and goldfish HSF1 (gHSF1) differ at amino acid residues 36 and 103. In hHSF1, both amino acid residues 36 and 103 are occupied by cysteine (C). However, in gHSF1, residue 36 is occupied by cysteine, whereas residue 103 is occupied by tyrosine. indicating that the disulfide bonds found in hHSF1 are not present in gHSF1 ^17^. A different residue, 103 was also found in walleye pollock which is a representative example of a Coldwater fish. Residue 103 in walleye pollock HSF1 (wHSF1) is occupied by phenylalanine. This also suggests that disulfide bonds may not be present in wHSF1. In this study, we focused on fish stress caused by water temperature and investigate the different HSF1, especially residue 103 in humans, goldfish, and walleye pollock and suggest how the different amino acid residual structures explain the different trimerization temperatures. (Fig 1)

**Figure 1.**
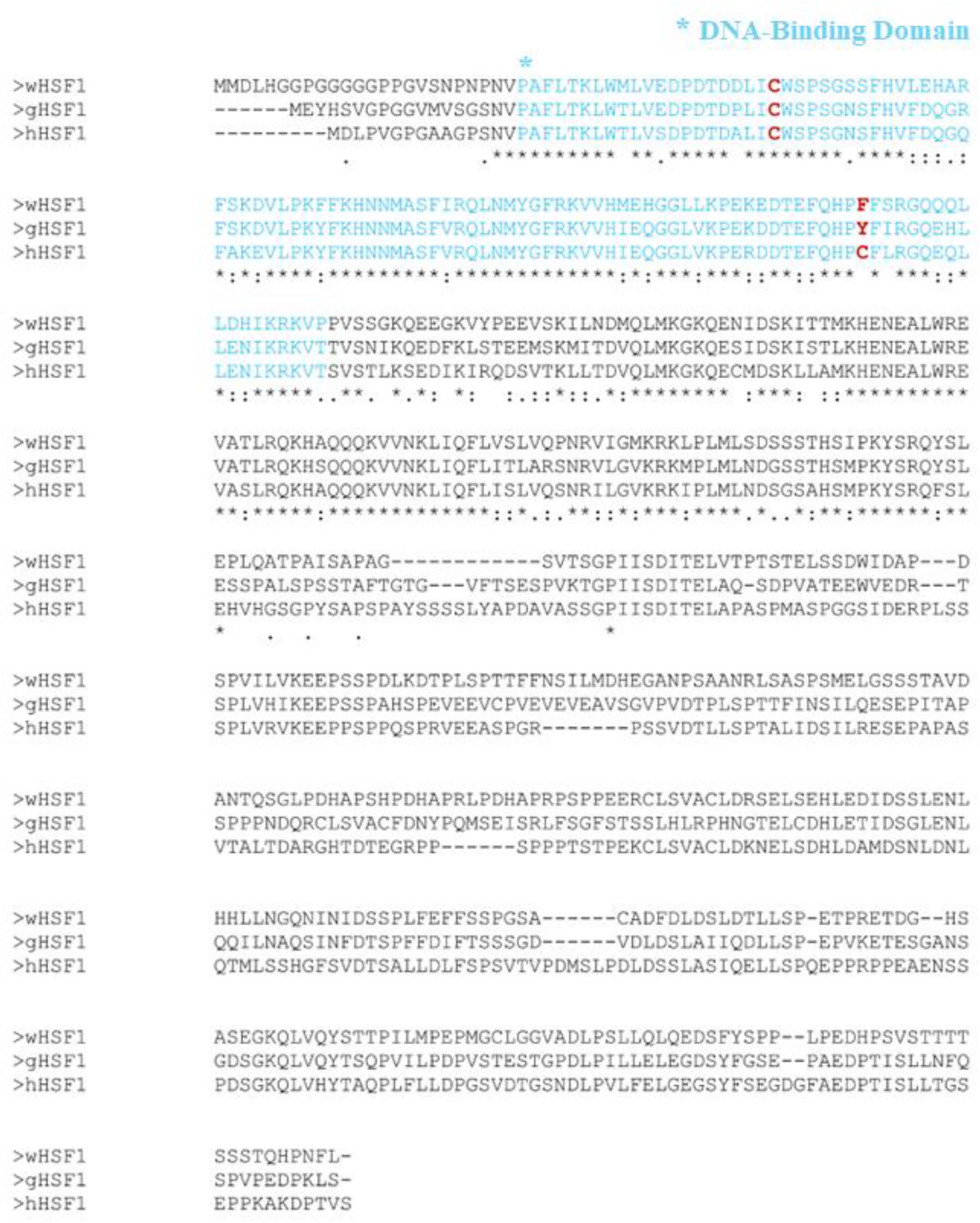
The alignment sequences of human, goldfish, and walleye pollock HSF1s.

## 2. Materials and methods

### 2.1 Isolation of HSF cDNA

Total RNA was isolated from the livers of walleye pollock in liquid nitrogen using an easyspin TM total RNA extraction kit (Invitrogen). The first-strand cDNA was synthesized from 5 µg of the total RNA with an oligo (dT) primer using the AMV reverse transcriptase (Promega). A polymerase chain reaction (PCR) was performed using the cDNA as a template, which contained 20 µM each of forward and reverse primers, Taq DNA polymerase, and 2.5 µM each of the deoxynucleotide triphosphates (dNTPs) (Bioneer, Korea). Primers were designed based on the conserved regions of Gadus morhua HSF1 (accession number XM_030346608). The PCR was performed for twenty-seven cycles of the following single cycle: 1 min at 95 °C for 30 s, 30 s at 55 °C, and 1 min at 70 °C. The PCR products were confirmed by electrophoresis, and the nucleotide sequences were determined using a 3730xl DNA analyzer (Macrogen).

### 2.2 Rapid amplification of cDNA 5’ and 3’ ends (RACE)

Full-length HSF1 and its sequences were obtained by the rapid amplification of cDNA ends using a SMART cDNA amplification kit (Clontech). Gene-specific primers (GSPs) were designed for the 5′-RACE and 3′-RACE procedures to obtain the full-length cDNA sequence of wHSF1. The 5′-RACE-PCR and 3′-RACE-PCR products were amplified using the GSPs and adapter primers included in the kit. The final PCR products were cloned into the TOPO PCR 2.1 vector (Invitrogen), and the sequences for full-length wHSF1, were determined. The sequences were deposited into GenBank with the accession number, MT350340.

### 2.3 Plasmids and vector construction

The expression vectors of wHSF1 were constructed by inserting full-length wHSF1 cDNA into a pET21b vector (Novagen). (hHSF1 and gHSF1 vector were provided by our laboratory). All the hHSF1 (accession number XM_937718), gHSF1 (accession number KJ145025), and wHSF1 residues were replaced with cysteine, tyrosine, phenylalanine, alanine, residues by PCR-mediated site-directed mutagenesis (Table 1). All plasmids were isolated from Escherichia coli JM109 cells and verified by DNA sequencing. ^18^.

**Table 1.**
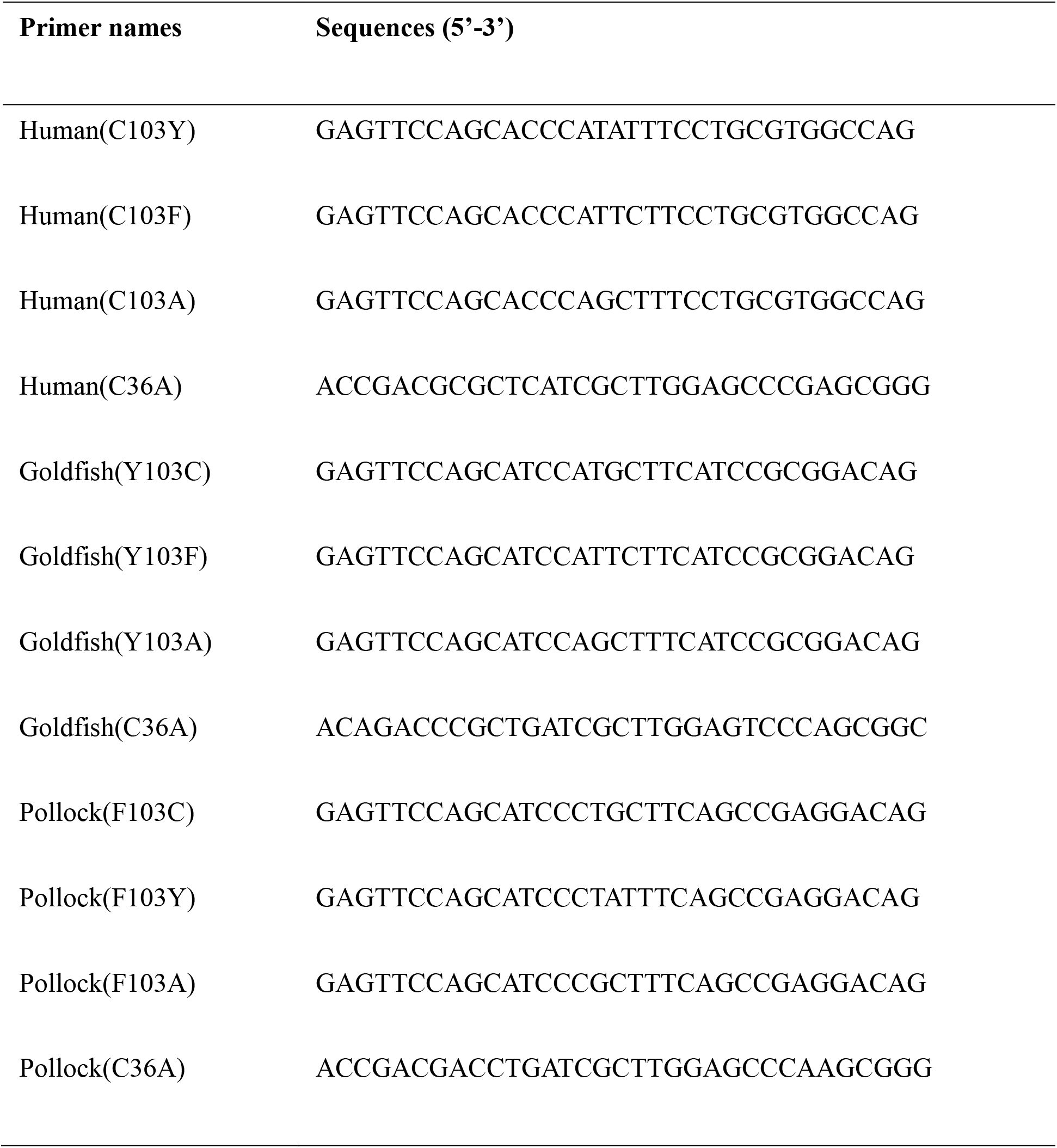
List of primers used for point mutation.

### 2.4 Expression and purification of protein

*E. coli* BL21(DE3) was transformed with full-length human, goldfish, and walleye pollock wildtype or their mutants. *E. coli* cells were cultured in a lysogeny broth (LB) medium to an OD_600_ of 0.5–0.6, and the proteins were induced by adding 1 mM of isopropyl-*β*-D-1-thiogalactopyranoside (IPTG) and keeping the sample at 20 °C for 24 h. The cells were collected by centrifugation and resuspended in a binding buffer containing 20 mM of Tris-HCl, 30 mM of imidazole, 200 mM of sodium chloride, and 1 mM of PMSF. After centrifugation (13,000 rpm, 30 min at 4 °C) of the crude lysate, the supernatant was purified using a His-Trap Ni column. ^19^.

### 2.5 HSF1 trimerization cross link assay

The trimerization activities of full-length hHSF1, gHSF1 and wHSF1 wildtype or its mutants were measured from crosslinking experiments. The purified proteins were incubated with 1 mM DTT and 1 mM ethylene glycol bis [succinimidyl succinate] (EGS) for 10 min on ice and mixed with five non-reducing sample buffers ^20^. Full-length hHSF1, gHSF1 and wHSF1 or its mutants were untreated or heat-activated for 30 min at different temperatures and then loaded onto a non-reducing 12% SDS-PAGE gel to determine the relative amounts of HSF1 monomers, dimers, and trimers. For western blotting, the loading on the non-reduction 12% SDS-PAGE was transferred to a 0.45 mm nitrocellulose membrane (Fisher). The membrane was blocked at 1 Tris at 0.05% Tween 20 and 3% BSA for 1 hour at room temperature. After washing once with a 1 Tris buffer containing 0.5% BSA, the membrane was incubated with an alkali phosphatase-bound chlorine-antibody (Sigma _A3562_), washed three times, and developed using a reinforced chemiluminescent substrate system (Promega).

### 2.6 Confirmation of the type of HSF1 binding (by heating after crosslink)

Two samples of hHSF1, gHSF1 and wHSF1 were prepared respectively, where crosslinking was completed (42, 37, and 20 °C) To determine covalent or non-covalent bonding, the temperature of one sample is the same as before, and the temperature of the other sample is raised to 55 °C for hHSF1, 42 °C for gHSF1, and 37 °C for wHSF1. Proceed with the western to determine the type of bonding of the two samples made. The method of Western blotting proceeds in the same way as before.

## 3. Results

### 3.1 The wildtype of HSF1 (human, goldfish, and walleye pollock) titration

First, we confirmed the temperature at which trimers were formed in humans, goldfish, and pollack. For hHSF1, the temperature was gradually increased and the temperature at which trimer was formed was measured through western blot. In the case of hHSF1, trimers were formed between 40-47°C, and the optimal temperature was confirmed to be 42°C. In gHSF1, trimers were formed between 30-37°C, and 37°C was confirmed to be the optimal temperature. Additionally, trimers were formed between 15-20°C in wHSF1, and 20°C was confirmed to be the optimal temperature.

### 3.2 The optimal temperature of wild-type HSF1s (human, goldfish, and walleye pollock)

For greater accuracy in the experiments, the temperatures of trimer formation were determined more accurately. The optimal temperatures of wild-type hHSF1, gHSF1, and wHSF1 trimers were 42, 37 and 20°C, respectively (Figure 3).

### 3.3 Trp-Floursence

Wild-type hHSF1, gHSF1 and wHSF1 were also examined by Trp-fluorescence spectroscopy that is used to identify the protein folding/unfolding ref. The wild-type hHSF1 after heat shock (42°C, 30 min) had a higher fluorescence intensity (FI) than that in its native state (Figure 2A). This result suggested that hHSF1 is a thermophilic protein, because hHSF1 could be induced to enter into a more folded (or more compact) state, but not into an unfolded one. Unlike the wild-type hHSF1 at 42°C, heat stimuli did not increase the FI value at 37 and 20°C Thus, these results demonstrated that hHSF1 forms a trimer at 42°C, and at lower temperatures of 37°C and 20°C, no trimer is formed and exists as a monomer (Figure 4A). And gHSF1 forms a trimer at 37°C, and trimer was not formed at 42°C and 20°C (Figure 4B). Also, wHSF1 forms a trimer at 20°C, and trimer was not formed at 42°C and 37°C (Figure 4C).

**Figure 2.**
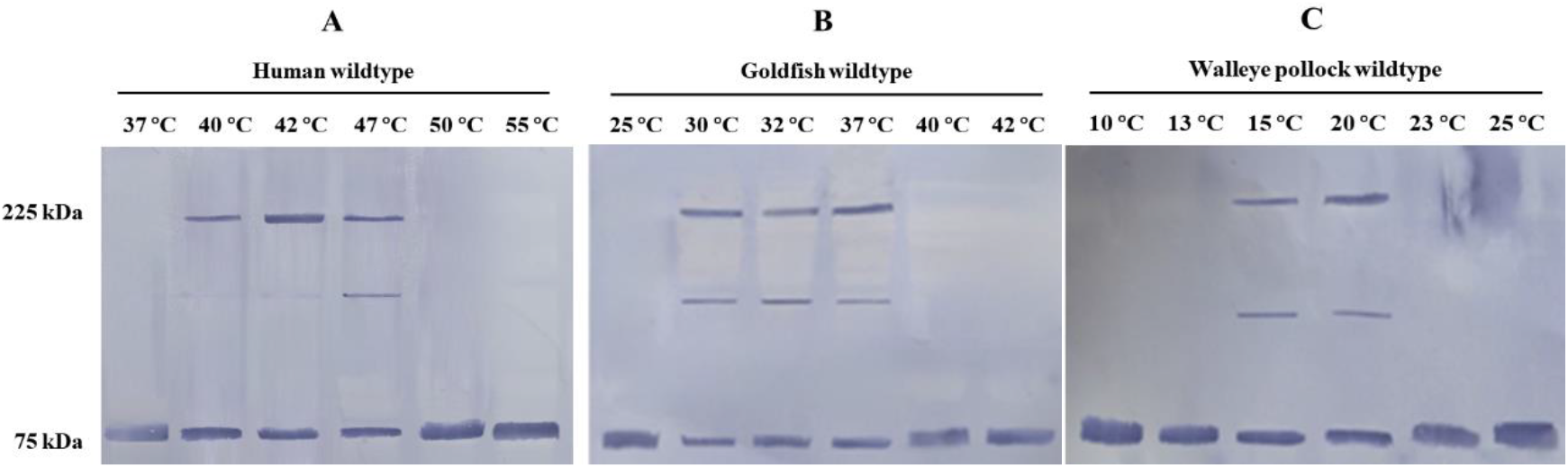
Trimerization of wild-type human, goldfish, and walleye pollock HSF1s. (A) is hHSF1, (B) is gHSF1, (C) is wHSF1.

**Figure 3.**
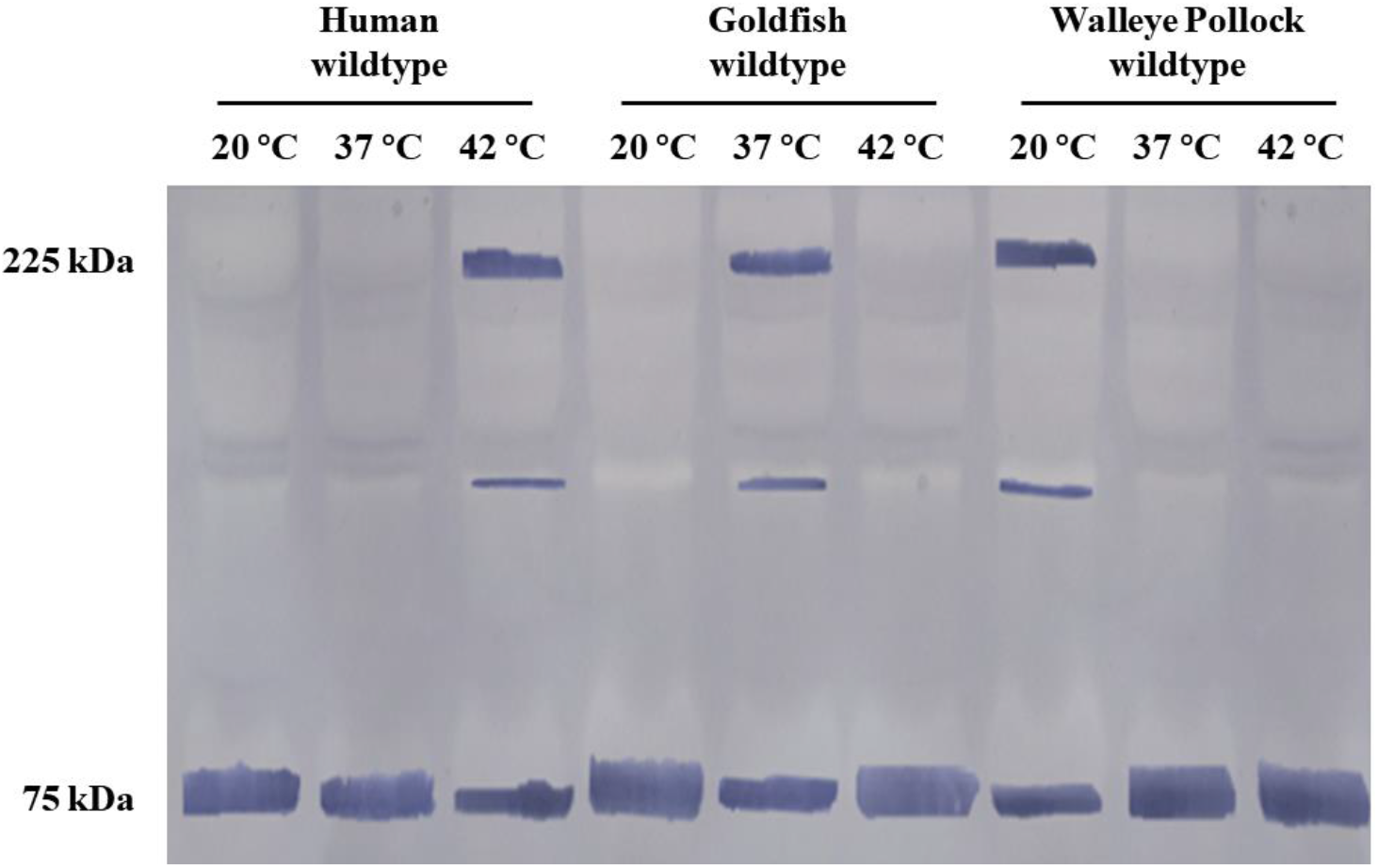
Optimal trimerization temperature of wild-type hHSF1, gHSF1, and wHSF1 in a temperature range of 20–42 °C by western blotting

**Figure 4.**
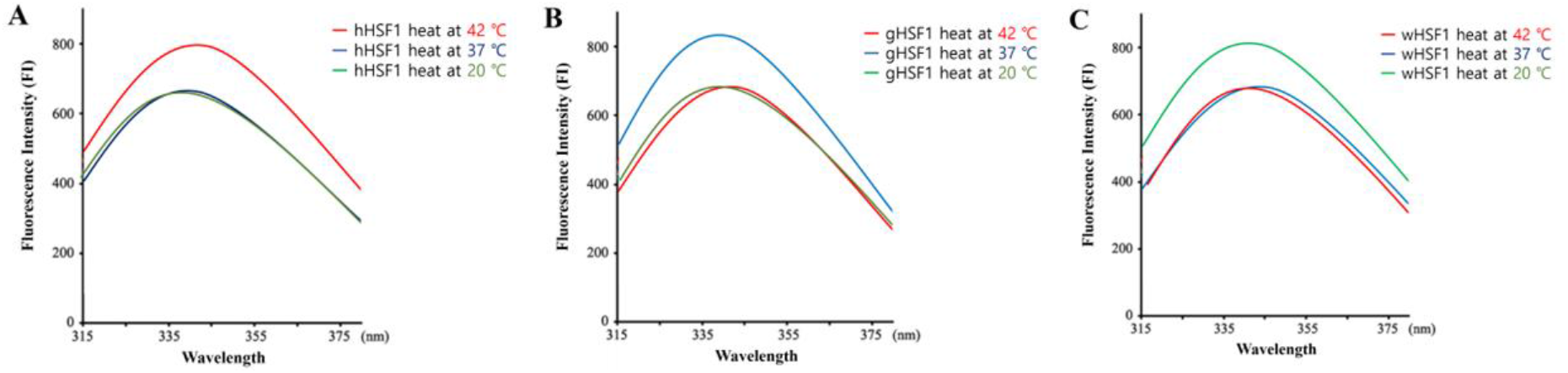
Trp-Fluorescence spectra of hHSF1, gHSF1 and wHSF1. Fluorescence samples (250μg/mL) were prepared in TGE buffer (50 mM Tris-HCl, 25% (v/v) glycerol, 1 mM EDTA, pH = 7.5). For the selective excitation of Trp amino acid, a wave length of 295 nm was used (the parameters were setup as λ_excitation_= 280 nm, λ_emission_=315-380 nm

### 3.3 The residue 103 mutation of HSF1 (human, goldfish, walleye pollock) with C, Y, F

Next, we determined whether the different amino acid residues at site 103 on the DNA binding domain were responsible for the different temperatures of trimer formation in hHSF1, gHSF1 and wHSF1. The wildtype HSF1 of each species was assigned a point mutation at site 103 with amino acid residues of the other two species. The results of the wildtype and the two mutations were compared using western blotting. hHSF1 formed a trimer at 42 °C, whereas the C103Y mutation formed a trimer at 37 °C and the C103F mutation formed a trimer at 20 °C (Figure 5A). In the goldfish specimens, gHSF1 formed a trimer at 37 °C, whereas the Y103C mutation formed a trimer at 42 °C and the Y103F mutation formed a trimer at 20 °C (Figure 5B). wHSF1 formed a trimer at 20 °C, whereas the F103C mutation formed a trimer at 42 °C and the F103Y mutation formed a trimer at 37 °C (Figure 5C). These results show that trimerization occurs at different temperatures depending on the amino acid residue at site 103, regardless of the wildtype HSF1 species.

**Figure 5.**
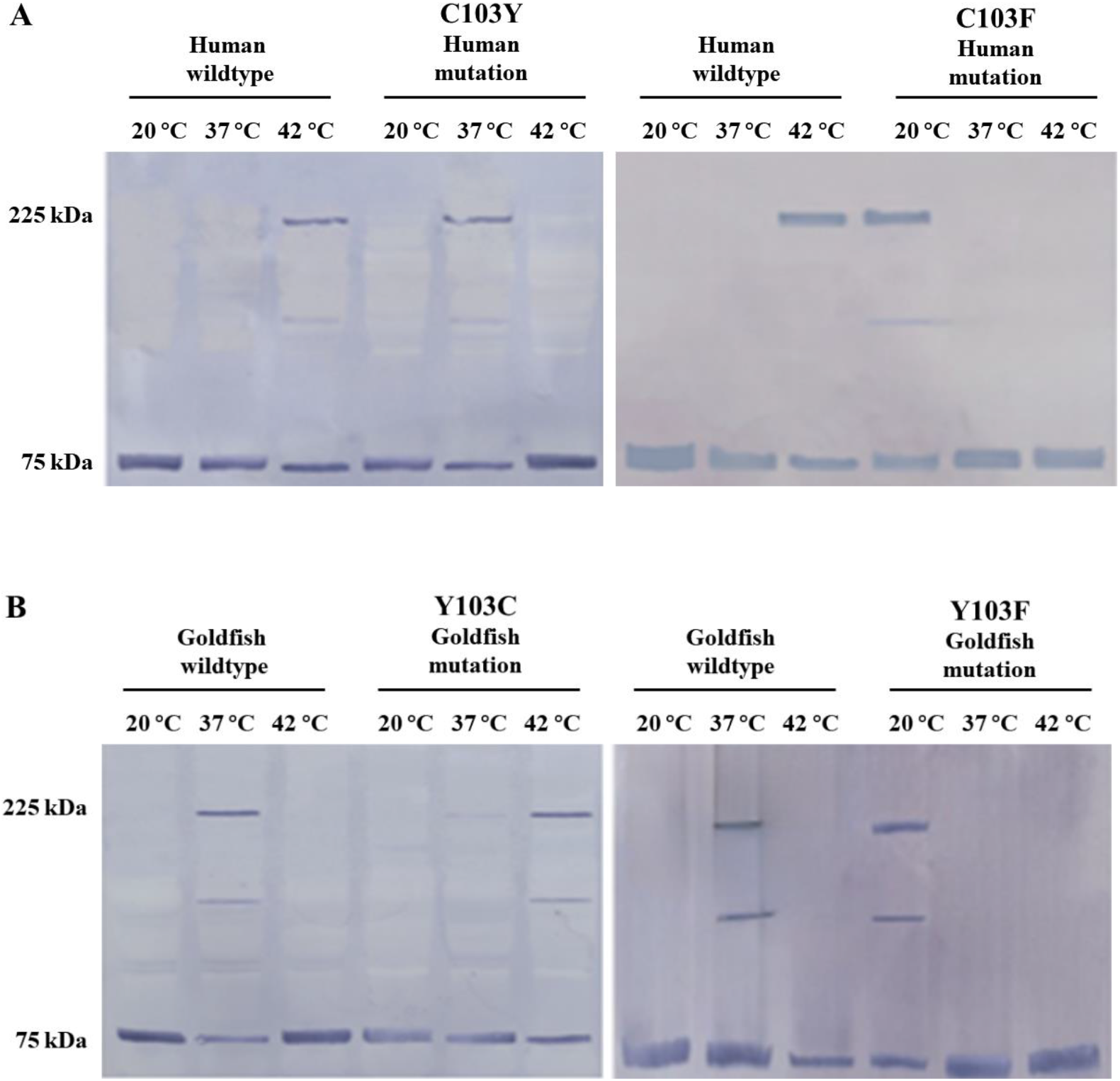
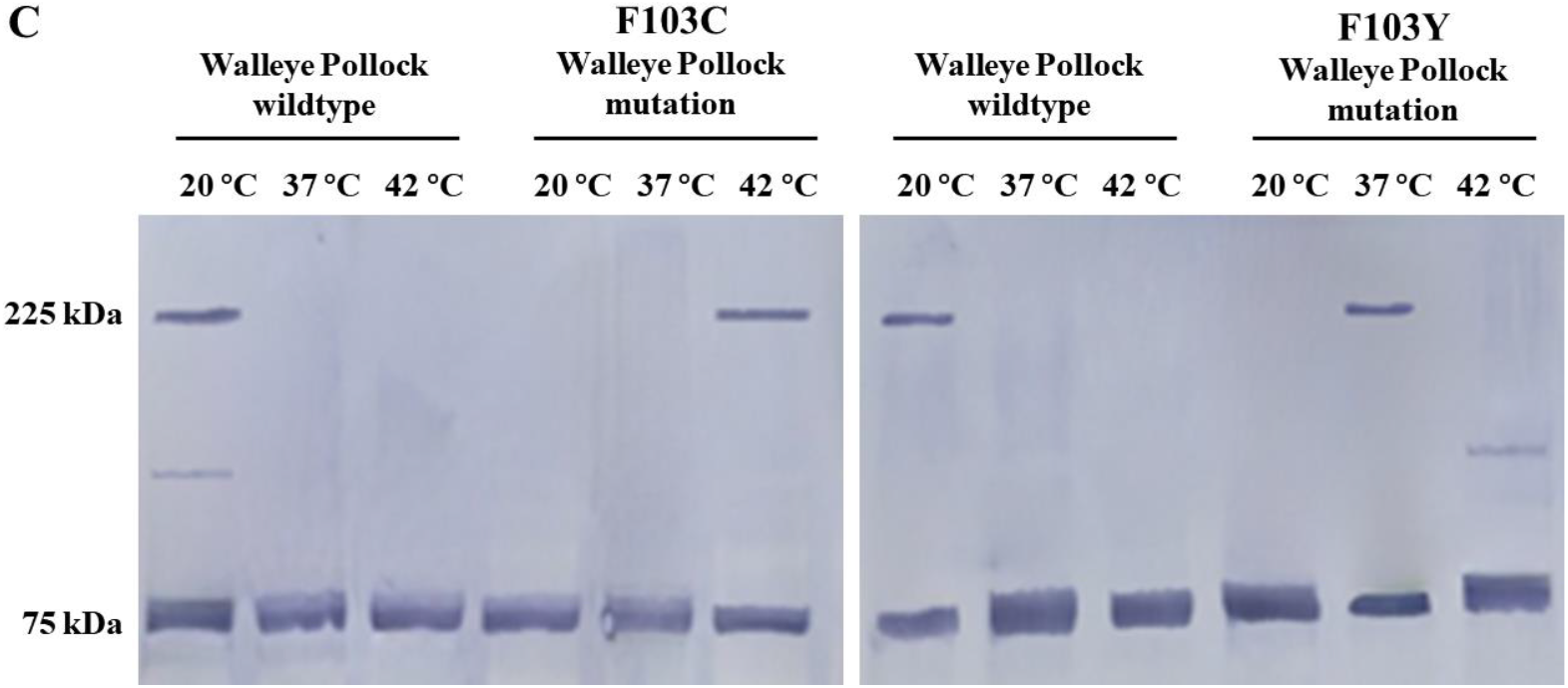
Trimerization of human, goldfish, and walleye pollock HSF1s and its mutants (residue at site 103) at 20–42 °C by western blotting: (A) human mutants: lanes 1–3 are wildtype control and lanes 4–6 are mutant; (B) goldfish mutants: lanes 1–3 are wildtype for control and lanes 4–6 are mutant; (C) walleye pollock mutants: lanes 1–3 are wildtype for control and lanes 4–6 are mutant.

### 3.4 The residue 103 mutation of HSF1 (human, goldfish, and walleye pollock) with A

A previous study showed that changing the amino acid residue at site 103 directly affected the trimerization temperature of wildtype HSF1s. The two residues at position 103 in goldfish and walleye pollock, tyrosine and phenylalanine, respectively, are aromatic amino acids with phenyl moiety. We wanted to confirm whether these amino acid residues are important for the trimerization of wildtype HSF1. The amino acid residue at site 103 was changed from the respective residues found in human, goldfish, and walleye pollock HSF1s to alanine, a nonaromatic amino acid. The results of the western blots showed that while wildtype HSF1s of each species formed trimers at their given temperatures, HSF1s with mutations to alanine at site 103 resulted in no trimerization in the temperature range of 20–42 °C (Figures 6A–C). These results indicate that trimerization does not occur when site 103 is occupied by a nonaromatic amino acid.

**Figure 6.**
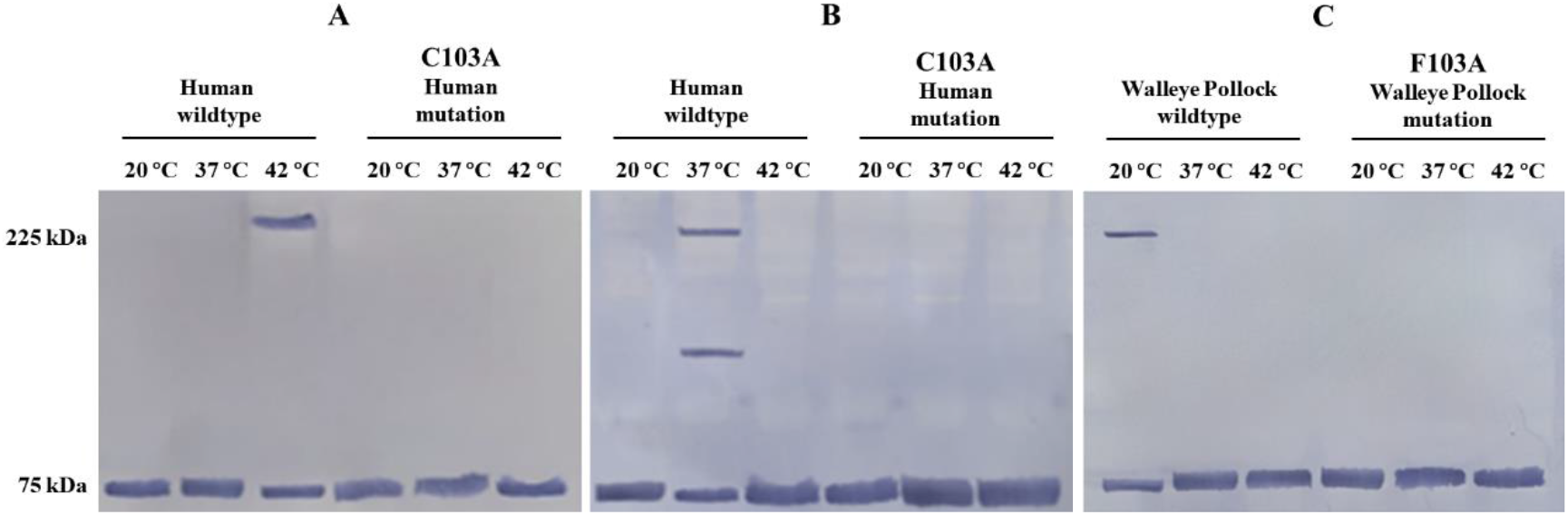
Trimerization of human, goldfish, and walleye pollock HSF1s and their mutants (residue at site 103) at 20–42 °C by western blotting: (A) human mutants: lanes 1–3 are wildtype control and lanes 4–6 are mutants; (B) goldfish mutants: lanes 1–3 are wildtype for control and lanes 4–6 are mutants; (C) walleye pollock mutants: lanes 1–3 are wildtype for control and lanes 4-6 are mutant.

### 3.5 The residue 36 mutation of HSF1 (human, goldfish, and walleye pollock) with C, Y, F

The previous experiments showed the importance of cysteine or aromatic compounds at site 103 for the trimerization of wildtype HSF1s. We wanted to find out if the presence of cysteine at site 36 for all three wildtype HSF1s were necessary for the trimerization to occur. The cysteine at site 36 for all three wildtype HSF1s were mutated to tyrosine and phenylalanine. The human HSF1s, with C36Y and C36F mutations, formed trimers at 37 °C and 20 °C respectively (Figure 5A). However, when goldfish and pollock HSF1s were treated with each mutation, trimerizations did not occur (Figure 5B, 5C). This showed that because human HSF1 had cysteine at site 103, trimerization was able to occur, albeit at different temperatures. However, in goldfish and pollock HSF1s, the lack of cysteines at both site 36 and site 103 resulted in no trimerization. This showed that at least one cysteine is required for the trimerization of HSF1s, and that it does not matter whether the cysteine is found at site 36 or 103.

### 3.6. The residue 36 mutation of HSF1 (human, goldfish, and walleye pollock) with A

Previous studies have shown the importance of cysteine or aromatic amino acids with phenyl moiety at site 103 in the trimerization of wild type HSF1s. The purpose of this experiment was to see if any direct bonds formed between cysteine 36 and residue alterations at 103 to produce a trimer, which, if proven true, would greatly enhance our understanding of the HSF1 trimer mechanism. Switching residue 36 of hHSF1, gHSF1, and wHSF1 to A (alanine) by the method of point mutation confirmed that no trimerization occurred in any of the human, goldfish, and walleye pollock samples (Figures 8A-C). These observations were in alignment with previous study that proved hHSF1 trimer is formed by the disulfide bond of 36 cysteine and 103 cysteine, that gHSF1 and wHSF1 also had 36 cysteine and 103 aromatic amino acids directly combine to form a trimer.

**Figure 7.**
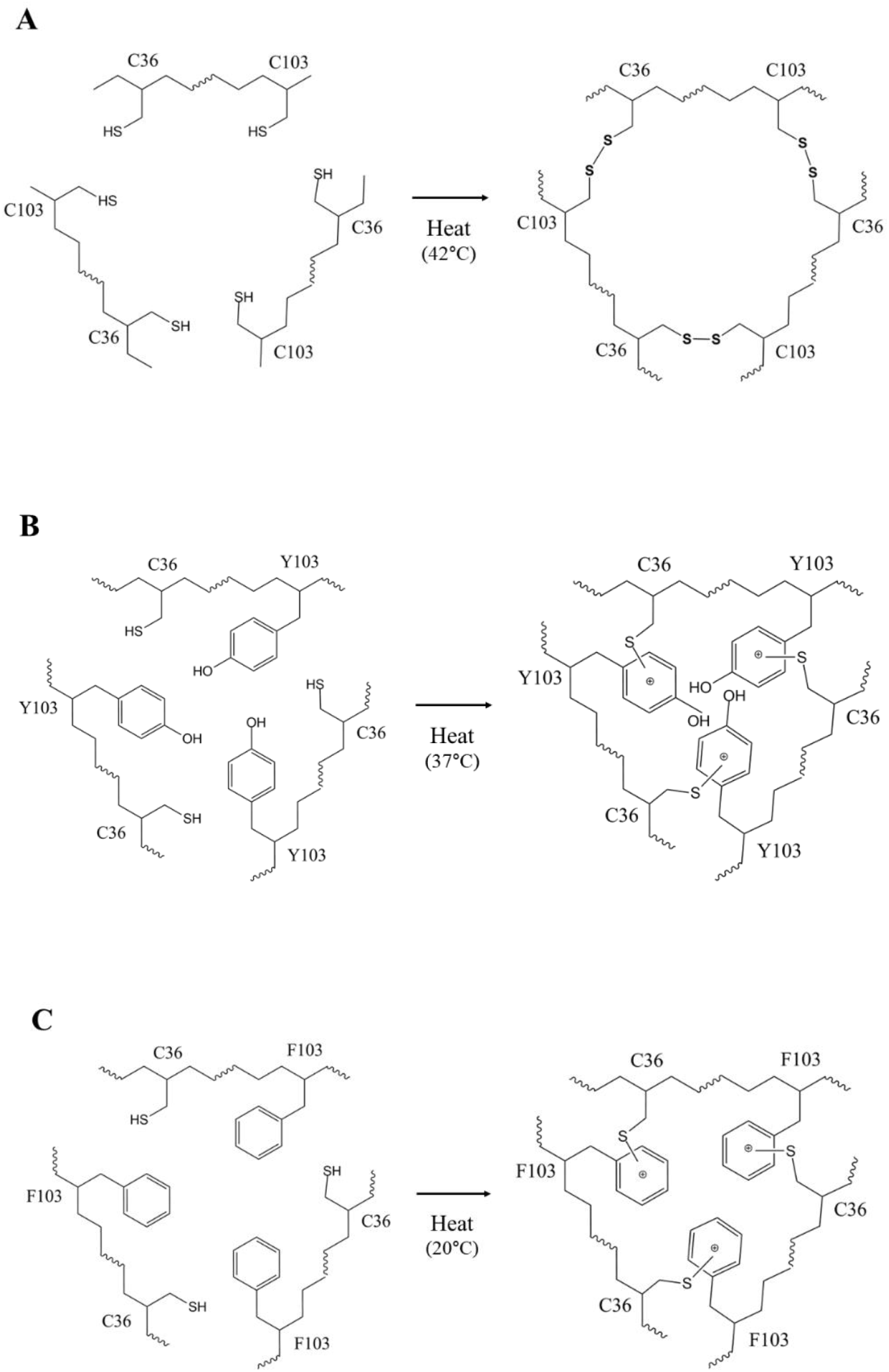
Trimerization of human, goldfish, and walleye pollock HSF1s and their mutants (residue at site 36) at 20–42 °C by western blotting: (A) human mutants: lanes 1–3 are wildtype for control and lanes 4–6 are mutants; (B) goldfish mutants: lanes 1–3 are wildtype for control and lanes 4–6 are mutants; (C) walleye pollock mutants: lanes 1–3 are wildtype for control and lanes 4-6 are mutant.

**Figure 8.**
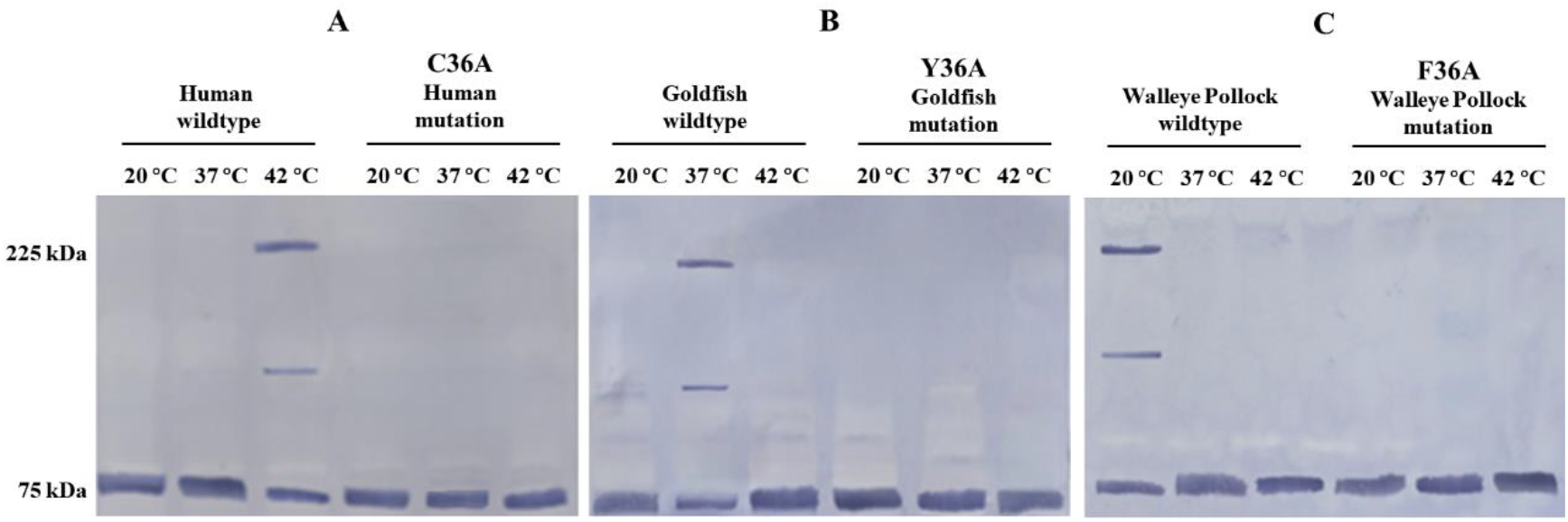
Trimerization of human, goldfish, and walleye pollock HSF1s and their mutants (residue at site 36) at 20–42 °C by western blotting: (A) human mutants: lanes 1–3 are wildtype for control and lanes 4–6 are mutants; (B) goldfish mutants: lanes 1–3 are wildtype for control and lanes 4–6 are mutants; (C) walleye pollock mutants: lanes 1–3 are wildtype for control and lanes 4-6 are mutant.

### 3.7 Confirmation of the type of bonds found at sites 36 and 103

The previous results have shown the trimerization of three different wildtype HSF1s at different temperatures. The two cysteines, as found in hHSF1, are known to form a disulfide covalent bond. With gHSF1 and wHSF1, the cysteine and the aromatic amino acids with phenyl moiety would form non-covalent bonds. We decided to test the strengths of these bonds by cross-linked trimerization at higher temperatures. The higher temperatures for each species (human: 55 °C, goldfish: 42 °C, and walleye pollock: 37 °C) were based on temperatures at which trimerization of respective HSF1s did not occur, even with the aid of cross-linking. Each wildtype HSF1s were cross-linked at their trimerization temperatures and then heated to the higher temperatures mentioned. The results were seen in western blots. In hHSF1, the cross-link between two cysteines were formed at 42 °C and temperature was raised to 55 °C before running a western blot. The trimerization still remained intact, showing that the covalent bond did not break (Figure 9A). For gHSF1 and wHSF1, the cross-link was formed at their respective trimerization temperatures, and the temperature was raised to 42 °C and 37 °C respectively before western blotting. The results showed that the non-covalent bonds did not survive the rise in temperature and trimerization did not remain intact (Figure 9B).

**Figure 9.**
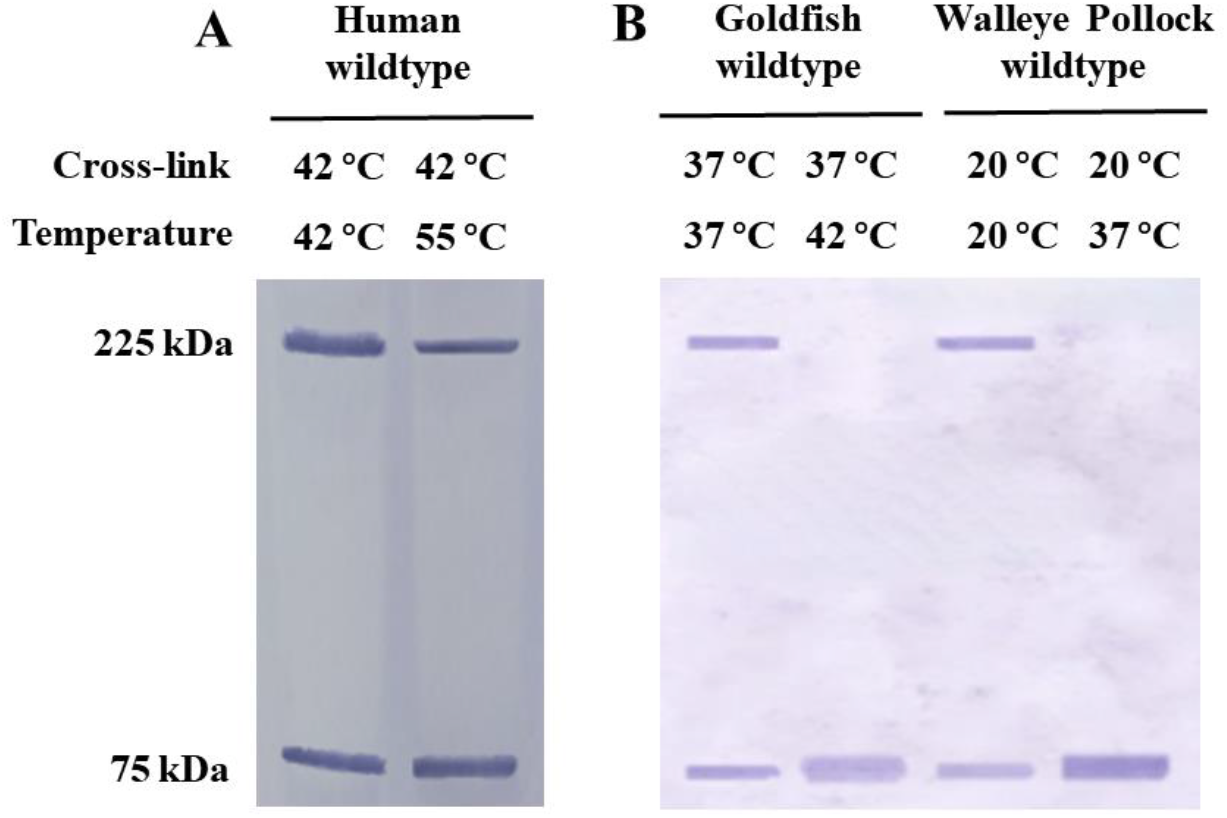
Western blotting of human, goldfish, and walleye pollock wildtype HSF1s to determine the types of bonds.

**Figure 10.**
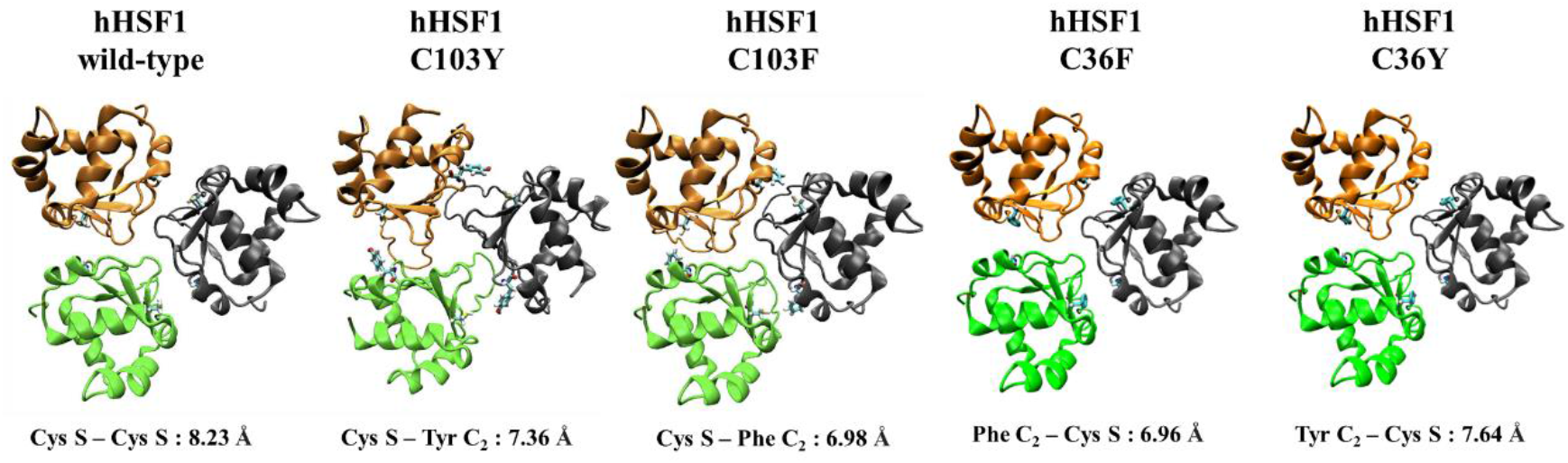

### 3.8 Modeling of Trimer Structures by Visual Molecular Dynamic

The initial structures of the HSF1 DNA binding domains (DBD) were predicted by AlphaFold2 (cite: https://www.nature.com/articles/s41586-021-03819-2), which produces the most probable structure from a given protein sequence. Five monomer structures were generated for each of the human, goldfish, and gadus DBDs. To predict the trimeric structure, we employed ROSIE (cite: https://journals.plos.org/plosone/article?id=10.1371/journal.pone.0063906), a protein modeling web server (https://rosie.rosettacommons.org/) based on the Rosetta (cite: https://www.annualreviews.org/doi/full/10.1146/annurev.biochem.77.062906.171838) framework. Using the symmetric docking protocol equipped on ROSIE, we could impose a symmetry constraint during the trimerization process, resulting in the generation of 400 symmetry-optimized trimeric structures for each monomeric structure. The distances between the residues of interest were calculated in each structure to identify the structures that could explain the experimental data. Mutations were introduced using the Chimera (cite: https://onlinelibrary.wiley.com/doi/10.1002/jcc.20084) software, generating alternativestructures with mutations at residue 22.

## 4. Discussion

All living organisms survive by adapting to the environment and temperature in which they live, whether animals on land or fish living in water. HSF1 studies in many species have continued. (human, mouse, chicken, goldfish, carp)^20–25^. The focus of this study was the trimerization of human, goldfish, and walleye pollock HSF1 at different temperatures and the mechanism of this trimerization. Previous study has shown that human HSF1 is characterized by two cysteines located in the DNA-binding domain, resulting in S-S bonds and a high trimerization temperature of about 42 °C ^12^. Previous studies have shown that goldfish HSF1 lacks S-S bonds and trimerization occurs at about 37 °C ^17^. We performed HSF1 trimerization assays on human, goldfish, and walleye pollock wildtype HSF1 samples based on previously developed methods and confirmed that trimers were formed at 42, 37, and 20 °C in the human, goldfish, and walleye pollock HSF1s, respectively. (Fig 2, 3)

The purpose of our study was to investigate the temperature and mechanism of trimerization formation by introducing point mutations in specific sequences. Previous studies have shown that in human HSF1, cysteine residues 36 and 103 form disulfide bonds at 42°C ^12^. In goldfish and walleye pollock HSF1s, while site 36 is occupied by cysteine, site 103 is occupied by tyrosine and phenylalanine, respectively. Considering that amino acids C, Y and F at residue 103 of these three species of HSF1 are factors in the trimer formation temperature, amino acid residue 103 of HSF1 of these three species was mutated. By introducing a point mutation of C103Y at site 103 to hHSF1, the temperature of trimerization decreased from 42 °C to 37 °C. In a similar experiment with the C103F mutation, the temperature was found to decrease from 42 °C to 20 °C. The temperature of trimerization was also tested for goldfish and walleye pollock, where their HSF1s were point-mutated at site 103 with the amino acid residues of the other two HSF1s. The experiments showed that, regardless of the species of HSF1, the trimerization temperature depends only on cysteine and the amino acid residue at site 103 (Fig. 5). This suggests that HSF1 residue 103 is a temperature-regulating factor for trimer formation.

A previous study showed that in hHSF1, when cysteine at site 103 was changed to serine, trimerization did not occur^15,12^. We conducted a similar experiment in which the cysteine at site 103 in all three HSF1s (human, goldfish, and walleye pollock) was converted to alanine (Fig 4). This mutated HSF1 did not undergo trimerization. In human HSF1, residue 36 is directly involved in trimer formation. In addition, in all three wildtype HSF1s, cysteine is present at site 36. We tested the importance of the cysteine at site 36 in trimerization. In a previous study, a study showed that trimers were not formed even when human HSF1 C1 was mutated to S. Residue 36 was mutated from cysteine to alanine in all three wildtype HSF1s. This mutated HSF1 also did not undergo trimerization (Fig 5). These results lead to the following conclusions: First, all three species, cysteine 36 is essential for trimerization to occur. Second, cysteine, tyrosine, and phenylalanine at residue 103 are not only HSF1 temperature regulatory factors but are also required for the trimerization mechanism of HSF1.

Based on the above evidence, we conducted several more experiments to predict the HSF1 mechanism of goldfish and walleye pollock. For gHSF1 and wHSF1, the cross-link was formed at their respective trimerization temperatures, and the temperature was raised to 42 °C and 37°C respectively before western blotting. The results showed that the non-covalent bonds did not survive the rise in temperature and trimerization did not remain intact (Fig 6B). These results showed that the covalent disulfide bonds were stronger than the noncovalent bonds formed between cysteine and aromatic amino acids with phenyl moiety of gHSF1 and wHSF1. Since there is only one C in the goldfish and walleye pollock HSF1 DNA binding domain, the type of trimer formation bond was first confirmed. As a result, it was confirmed to be a non-covalent bond, and it can be proposed to be a non-covalent bond between C-Y in the case of goldfish and a non-covalent bond between C-F in the case of walleye pollock. This provides further evidence that the trimerization mechanism is related to either the disulfide bond between two cysteine residues (S-S) or a non-covalent bond between cysteine and aromatic amino acids with phenyl moiety. In previous studies, it was confirmed that aromatic amino acid phenyl moiety and -S of cysteine can form a non-covalent bond ^26–27^. Based on this, the trimmer structure of hHSF1, gHSF1, and wHSF1 was schematically illustrated (Fig 7).

**Figure 7.**
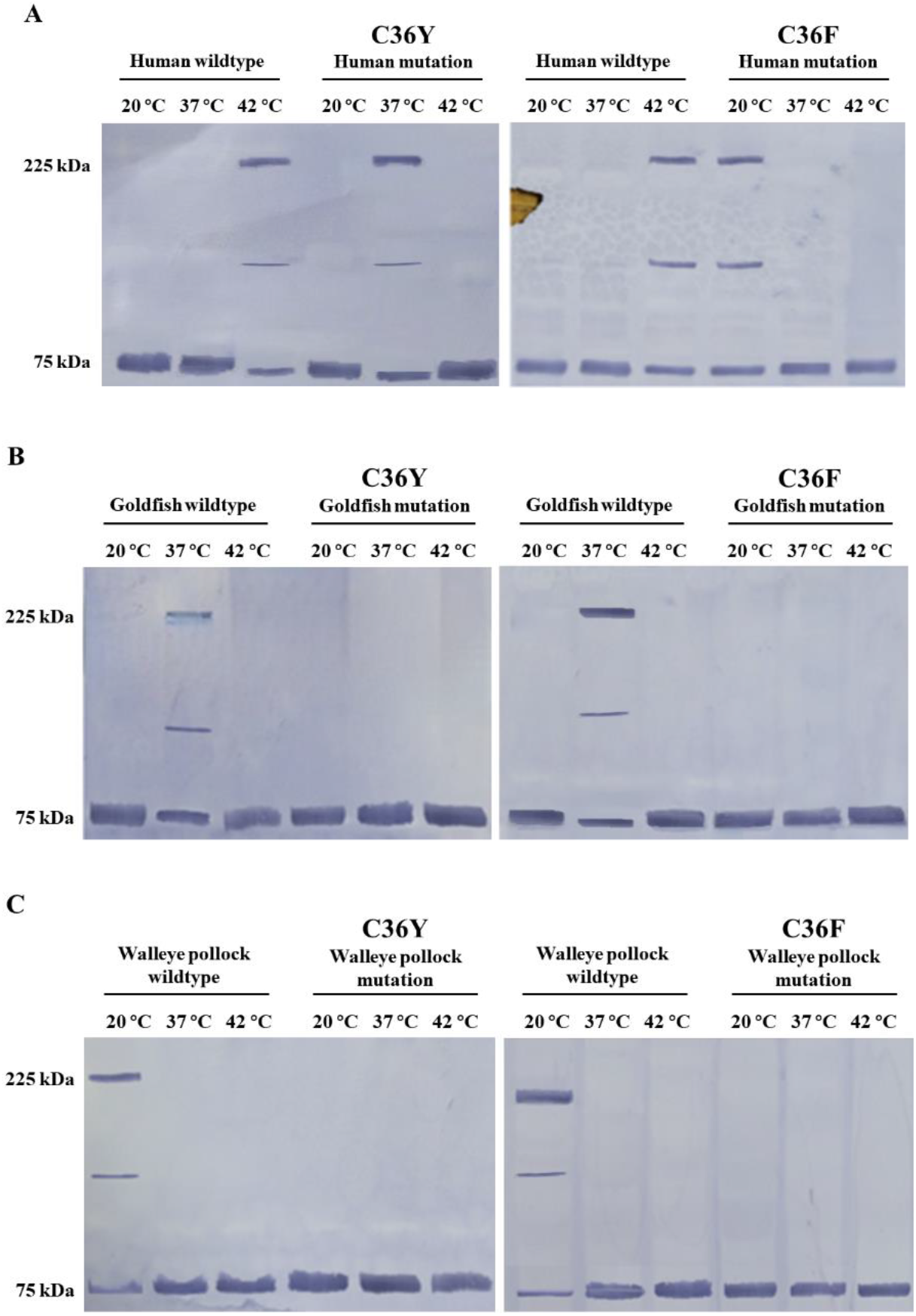
Diagrammatic illustration of intermolecular trimer interactions in human, goldfish, and walleye pollock wildtype HSF1: (A) hHSF1; (B) gHSF1; and (C) wHSF1.

The cumulative results suggest that the temperature of trimerization is correlated with the strength of the disulfide and non-covalent bonds in the wildtype HSF1s. The two cysteines in sites 36 and 103 in wildtype human HSF1 have been shown to lead to the formation of disulfide bonds, which helps with the formation of trimers at 42 °C. In goldfish and walleye pollock HSF1s, site 103 is occupied by tyrosine and phenylalanine, respectively, and the temperatures of trimerization for the goldfish and walleye pollock HSF1s are 37 °C and 20 °C, respectively. The amino acids tyrosine and phenylalanine are structurally similar in that they both contain aromatic rings. The key difference is that tyrosine contains a hydroxyl group (-OH) on its aromatic ring, whereas phenylalanine does not. This could explain why these aromatic amino acids undergo trimerization with cysteine at different temperatures. The bond between cysteine sulfide and the hydroxyl group of tyrosine could be stronger than that between cysteine sulfide and the phenylalanine aromatic ring. Another conclusion from these results is that the disulfide covalent bond in human HSF1 is strong enough to form trimers at 42 °C, whereas the reason for the trimerization temperatures of 37 °C and 20 °C for goldfish and walleye pollock, respectively, is the weaker non-covalent bonds between cysteine and the aromatic residues.

These results confirm that the temperature at which HSF1 trimerization occurs is closely related to the temperature of the environment. DNA binding activity of human HSF1 in different human cells is rather low below 40°C and strongly increased above 42°C ^28–30^. Other studies have shown that for goldfish, more than half of their population died when the surrounding water temperature exceeded 40 °C ^31^. For walleye pollock, this temperature was 19∼20 °C ^32–33^. This indicates that the HSF1 trimer formation. Comparing the HSF1 sequences of fish, Cold-water fish such as walleye pollock, Eleginops maclovinus, Chionodraco rastropinosus, and Pleuronectes platessa have residues of F 103 ^34–35^. While, Fish that are known to reside in warm water, such as goldfish, carp, catfish, and zebrafish, have a residue of Y 103 ^36,25,37^. This indicate that their HSF1 residue amino acid 103 is related to the water temperature environment in which they live. It also suggests that the HSF1 trimer mechanism is also related to their water temperature and survival. The results of these HSF1 experiments in goldfish and walleye pollock are thought to be useful not only for biochemical concepts but also for their aquaculture aspects. Lastly, in order to more accurately understand the HSF1 mechanism of goldfish and walleye pollock, our laboratory is conducting various HSF1 modeling using VMD (Visual molecular Dynamics).

## 5. Conclusion

The trimerization temperature of HSF1s from humans, goldfish, and walleye pollock was determined for different residues at site 103. For temperate fish, such as goldfish, residue 103 is Y, whereas it is F for Coldwater fish, such as walleye pollock. Goldfish and walleye pollock can form trimers owing to non-covalent bonds between cysteine and other aromatic amino acids with phenyl moiety acids, unlike humans, who form trimers with S-S bonds. Finally, the binding strength of trimerization determines the trimer formation temperature.

## Accession Codes

Human heat shock transcription factor 1, hHSF1 (accession number: XM_937718), Goldfish heat shock transcription factor 1, gHSF1 (accession number: KJ145025) and Walleye pollock heat shock transcription factor 1, wHSF1 (accession number: MT350340)

## Author contributions

Chang-Ju Lee: Writing-original draft, Methodology. Bo-Hee Choi: Methodology, Investigation, Software. So-Sun Kim: Methodology, Resources. David Nham-Joon Kim: Methodology, Investigation. Jeong-Mo Choi: Project advice. Young-Shang Park: Project Administration, Supervision. Jang-Su Park: Supervision.

All authors have read and agreed to the published version of the manuscript

## Notes

Conflicts of Interest: The authors declare no conflict of interest.

## Acknowledgments

This study was funded by the National Institute of Fisheries Science, Ministry of Oceans and Fisheries, Korea, Grant number: R2023062

## Notes

### Competing Interest Statement

The authors have declared no competing interest.

## References

1. Madeira D.; Narciso L.; Cabral H.; Vinagre C.; Diniz M. Influence of temperature in thermal and oxidative stress responses in estuarine fish. Comparative Biochemistry and Physiology Part A: Molecular & Integrative Physiology. 2013, 166, 237–43.

2. Mayr M.; Kiechl S.; Willeit J.; Wick G.; Xu Q. Associations of antibodies to Chlamydia pneumoniae, Helicobacter pylori, and cytomegalovirus with immune reactions to heat-shock protein 60 and carotid or femoral atherosclerosis. Circulation. 2000, 102, 833–9.

3. Airaksinen S.; Råbergh CM.; Lahti A.; Kaatrasalo A.; Sistonen L.; Nikinmaa M. Stressor-dependent regulation of the heat shock response in zebrafish, Danio rerio. Comparative Biochemistry and Physiology Part A: Molecular & Integrative Physiology. 2003, 134, 839–46.

4. Misra S.; Zafarullah M.; Price-Haughey J.; Gedamu L. Analysis of stress-induced gene expression in fish cell lines exposed to heavy metals and heat shock. Biochimica et Biophysica Acta (BBA)-Gene Structure and Expression. 1989, 1007, 325–33.

5. Åkerfelt M.; Morimoto RI.; Sistonen L. Heat shock factors: integrators of cell stress, development and lifespan. Nature reviews Molecular cell biology. 2010, 11, 545–55.

6. Archana P.; Aleena J.; Pragna P.; Vidya M.; Niyas A.; Bagath M. Role of heat shock proteins in livestock adaptation to heat stress. J Dairy Vet Anim Res. 2017, 5, 00127.

7. Feder ME.; Hofmann GE. Heat-shock proteins, molecular chaperones, and the stress response: evolutionary and ecological physiology. Annual review of physiology. 1999, 61, 243–82.

8. Vihervaara A.; Sistonen L. HSF1 at a glance. Journal of cell science. 2014, 127, 261–6.

9. Morimoto RI.; Tissières A.; Georgopoulos C. Stress proteins in biology and medicine. 1990.

10. Fujimoto M.; Nakai A. The heat shock factor family and adaptation to proteotoxic stress. The FEBS journal. 2010, 277, 4112–25.

11. Pirkkala L.; Nykänen P.; Sistonen L. Roles of the heat shock transcription factors in regulation of the heat shock response and beyond. The FASEB Journal. 2001, 15, 1118–31.

12. Lu M.; Kim H-E.; Li C-R.; Kim S.; Kwak I-J.; Lee Y-J.;, et al. Two distinct disulfide bonds formed in human heat shock transcription factor 1 act in opposition to regulate its DNA binding activity. Biochemistry. 2008, 47, 6007–15.

13. Wu C. Heat shock transcription factors: structure and regulation. Annual review of cell and developmental biology. 1995, 11, 441–69.

14. Zuo J.; Baler R.; Dahl G.; Voellmy R. Activation of the DNA-binding ability of human heat shock transcription factor 1 may involve the transition from an intramolecular to an intermolecular triple-stranded coiled-coil structure. Molecular and cellular biology. 1994, 14, 7557–68.

15. Ahn S-G.; Thiele DJ. Redox regulation of mammalian heat shock factor 1 is essential for Hsp gene activation and protection from stress. Genes & development. 2003, 17, 516–28.

16. Lu M.; Lee Y-J.; Park S-M.; Kang HS.; Kang SW.; Kim S.; Park J-S. Aromatic-participant interactions are essential for disulfide-bond-based trimerization in human heat shock transcription factor 1. Biochemistry. 2009, 48, 3795–7.

17. Kim S-S.; Chang Z.; Park J-S. Identification tissue distribution and characterization of two heat shock factors (HSFs) in goldfish (Carassius auratus). Fish & Shellfish Immunology. 2015, 43, 375–86.

18. Zheng L.; Baumann U.; Reymond J-L. An efficient one-step site-directed and site-saturation mutagenesis protocol. Nucleic acids research. 2004, 32, e115-e.

19. Soncin F.; Prevelige R.; Calderwood SK. Expression and purification of human heat-shock transcription factor 1. Protein expression and purification. 1997, 9, 27–32.

20. Kim S-S.; So J-H.; Park J-S. Comparison of Thermal Stress Induced Heat Shock Factor 1 (HSF1) in Goldfish and Mouse Hepatocyte Cultures. Life Science Journal. 2016, 26, 1360–6.

21. Rabindran SK.; Giorgi G.; Clos J.; Wu C. Molecular cloning and expression of a human heat shock factor, HSF1. Proceedings of the National Academy of Sciences. 1991, 88, 6906–10.

22. Manalo DJ.; Lin Z.; Liu AY-C. Redox-dependent regulation of the conformation and function of human heat shock factor 1. Biochemistry. 2002, 41, 2580–8.

23. Yan L-J.; Rajasekaran NS.; Sathyanarayanan S.; Benjamin IJ. Mouse HSF1 disruption perturbs redox state and increases mitochondrial oxidative stress in kidney. Antioxidants & redox signaling. 2005, 7, 465–71.

24. Nakai A.; Morimoto RI. Characterization of a novel chicken heat shock transcription factor, heat shock factor 3, suggests a new regulatory pathway. Molecular and Cellular Biology. 1993.

25. Yang X.; Gao Y.; Zhao M.; Wang X.; Zhou H.; Zhang A. Cloning and identification of grass carp transcription factor HSF1 and its characterization involving the production of fish HSP70. Fish physiology and biochemistry. 2020, 46, 1933–45.

26. Martinie RJ.; Godakumbura PI.; Porter EG.; Divakaran A.; Burkhart BJ.; Wertz JT.; Benson DE. Identifying proteins that can form tyrosine-cysteine crosslinks. Metallomics. 2012, 4, 1037–42.

27. Nauser T.; Casi G.; Koppenol WH.; Schöneich C. Intramolecular addition of cysteine thiyl radicals to phenylalanine in peptides: formation of cyclohexadienyl type radicals. Chemical communications. 2005, 3400–2.

28. Abravaya K.; Phillips B.; Morimoto RI. Attenuation of the heat shock response in HeLa cells is mediated by the release of bound heat shock transcription factor and is modulated by changes in growth and in heat shock temperatures. Genes & development. 1991, 5, 2117–27.

29. Baler R.; Dahl G.; Voellmy R. Activation of human heat shock genes is accompanied by oligomerization, modification, and rapid translocation of heat shock transcription factor HSF1. Molecular and cellular biology. 1993.

30. Mosser DD.; Kotzbauer PT.; Sarge KD.; Morimoto RI. In vitro activation of heat shock transcription factor DNA-binding by calcium and biochemical conditions that affect protein conformation. Proceedings of the National Academy of Sciences. 1990, 87, 3748–52.

31. Ford T.; Beitinger TL. Temperature tolerance in the goldfish, Carassius auratus. Journal of Thermal Biology. 2005, 30, 147–52.

32. Pérez-Casanova J.; Afonso L.; Johnson S.; Currie S.; Gamperl A. The stress and metabolic responses of juvenile Atlantic cod Gadus morhua L. to an acute thermal challenge. Journal of Fish Biology. 2008, 72, 899–916.

33. Kim S-S.; Lee C-J.; Yoo H-K.; Choi J.; Byun S-G.; Kim W-J. Effect of water temperature on walleye pollock (Gadus chalcogrammus) embryos, larvae and juveniles: Survival, HSP70 expression, and physiological responses. Aquaculture. 2022, 554, 738136.

34. Cha IS.; Kwon J.; Park SB.; Jang HB.; Nho SW.; Kim YK.;, et al. Heat shock protein profiles on the protein and gene expression levels in olive flounder kidney infected with Streptococcus parauberis. Fish & shellfish immunology. 2013, 34, 1455–62.

35. Daane JM.; Detrich III HW. Adaptations and diversity of Antarctic fishes: a genomic perspective. Annual Review of Animal Biosciences. 2022, 10, 39–62.

36. Dalvi RS.; Pal AK.; Tiwari LR.; Baruah K. Influence of acclimation temperature on the induction of heat-shock protein 70 in the catfish Horabagrus brachysoma (Günther). Fish Physiology and Biochemistry. 2012, 38, 919–27.

37. Rabergh C.; Airaksinen S.; Soitamo A.; Bjorklund H.; Johansson T.; Nikinmaa M.; Sistonen L. Tissue-specific expression of zebrafish (Danio rerio) heat shock factor 1 mRNAs in response to heat stress. Journal of Experimental Biology. 2000, 203, 1817–24.

